# Transmembrane helices 5 and 12 control transport dynamics, substrate affinity and specificity in the elevator-type UapA transporter

**DOI:** 10.1101/2022.04.04.486963

**Authors:** Dimitris Dimakis, Yiannis Pyrris, George Diallinas

## Abstract

An increasing number of solute transporters have been shown to function with the so-called *sliding-elevator* mechanism. Despite structural and functional differences all elevator-type transporters use a common mechanism of substrate translocation via reversible movements of a mobile core domain (the *elevator*) hosting the substrate binding site along a rigid scaffold domain stably anchored in the plasma membrane via homodimerization. One of the best studied elevator transporters is the UapA uric acid-xanthine/H^+^ symporter of the filamentous fungus *Aspergillus nidulans*. Here, we present a novel genetic analysis for deciphering the role of transmembrane segments (TMS) 5 and 12 in UapA transport function. We show that specific residues in both TMS5 and TMS12 control, negatively or positively, the dynamics of transport, but also substrate binding affinity and specificity. More specifically, mutations in TMS5 can lead to increased rate of transport, but also to an inactive transporter due to high-affinity substrate-trapping, whereas mutations in TMS12 lead to apparently uncontrolled sliding, and thus broadened specificity and UapA-mediated purine toxicity. Our findings shed new light on how elevator transporters function or how their transport characteristics might be altered genetically or have been modified in the course of evolution.

## Introduction

Transporters are transmembrane proteins that mediate the selective translocation of nutrients, metabolites or drugs across cellular membranes. Their activity is essential for cell survival, division and differentiation and consequently for the life of all organisms. Thus transporter malfunction is associated with several diseases, such as cystic fibrosis, diabetes or neurodegeneration. Transporters, despite their structural and functional differences, alternate in structurally distinct conformations during transport: outward-facing open (substrate reception), outward-facing occluded (substrate oriented in the major binding site and closure of outer gate), fully occluded (substrate stabilized in a major binding site), inward-facing occluded (substrate binding induces inward conformation) and inward-facing open (inner gate opens and substrate released) (Penmatsa et al, 2014; Simmons et al., 2014; Coincon et al,. 2016 Drew and Boudker, 2016). Based on their structure, mechanism of function and evolutionary origin, most transporters fall into three distinct groups, known as the Major Facilitator Superfamily (MFS), the Amino Acid-Polyamine-Organocation (APC) superfamily, and a more structurally diverse group of transporters operating by the so-called sliding elevator mechanism of transport, which necessitates functional dimerization (Saier, 2000; Drew and Boudker, 2016).

Elevator type transporters (symporters and antiporters or exchangers) possess a motile substrate binding site included in a core or elevator domain that moves through the membrane as a rigid body, translocating the substrate(s) from one side of the membrane to the other. The movement of the core/elevator domain takes place along a relatively rigid scaffold domain, anchored in the PM via homodimerization and specific interactions with lipids (Lee et al., 2013; Alguel et al., 2016; Byrne 2017; Garaeva and Slotboom, 2020; Pyle et al., 2018; Diallinas 2021). The basic aspects of this mechanism of transport have been deciphered by comparing outward- and inward-facing cryo-electron microscopy (EM) and X-ray crystallography structures of several elevator type transporters (Lee et al, 2013; Alguel et al., 2016; Geertsma et al., 2015; Coincon et al., 2016; Thurtle-Schmidt et al., 2016; Ficici et al., 2017; Yu et al., 2017). Structural studies on elevator-type transporters have been complemented and supported by extensive mutational and functional *in vivo* studies in UapA, a uric acid/xanthine transporter of the filamentous fungus *Aspergillus nidulans* (Koukaki et al. 2005; Vlanti et al., 2006; Pantazopoulou and Diallinas, 2006; Papageorgiou et al. 2008; Kosti et al. 2010; Amilllis et al. 2011; Kourkoulou et al., 2019; 2021). In particular, the genetic studies in UapA have contributed in understanding how substrate specificity is determined, an issue hard to approach solely via structural studies. Most interestingly, specificity mutations enlarging the UapA binding to all purines and uracil do not modify residues of the substrate binding site (i.e. in TMS3, TMS8 or TMS10), as might have been expected. Instead, they concern residues in TMS12, TMS13 or TMS14, which are helices of the rather rigid scaffold/dimerization domain. Other specificity mutations concerned residues in flexible loops linking the core/elevator and the scaffold domains. On the other hand, mutations in residues involved in direct substrate co-ordination and transport, located in TMS3, TMS8 and TMS10 of the movable elevator, alter the transport kinetics of UapA in respect to its native substrates, but generally do not modify specificity for nucleobases other than physiological substrates (for a recent review see Diallinas, 2016).

Homology modeling of the outward-facing conformation and comparison with inward-facing crystal structure of UapA provided an explanation on how specificity can be modified by affecting the dynamics of sliding of the core/elevator domain (Diallinas, 2021). In the outward-facing conformation and in the absence of substrate the core/elevator domain seems to be “locked” by tight interactions with specific residues in the scaffold domain and in particular with TMS12 and TMS14. Protonation of the substrate binding site and subsequent substrate binding seems to ‘unlock’ the sliding of the elevator by altering the polar character of the substrate binding site, which can now slide towards the cytoplasmic side of the PM along the interface with the scaffold domain, and in particular along TMS12 and TMS5. Through this model, specificity mutations can be rationalized as amino acid modifications that somehow facilitate sliding or uncouple it from the requirement of high-affinity substrate binding. As a consequence, any purine or other structurally similar solute that has access and fits into the binding site cavity, even with low affinity, can be transported by UapA.

In this work we investigate the mechanism of UapA transport function by designing and functionally analyzing novel mutations in TMS5 and TMS12, which form the sliding interface of the elevator. We present evidence that specific residues in both TMS5 and TMS12 control negatively or positively the dynamics of sliding of the elevator and thus also modify transport kinetics and specificity of UapA.

## Results and Discussion

### TMS5 mutations

The crystal structure and modeling of UapA has shown that TMS5 and TMS12 form the interface along which the core/elevator domain slides and mediates substrate/H^+^ symport. Specificity mutations have been isolated via unbiased genetic screens along TMS12 (Papageorgiou et al. 2008; Kosti et al. 2010), which is in line with the proposed model on how the dynamics of sliding might lead to enlarged UapA specificity (Diallinas, 2021). Curiously, no specificity mutation has been isolated in TMS5. This seems rather unexpected considering also that TMS5 is the most highly conserved TMS in UapA (**Figure 1A**). The single known mutation in TMS5 is L234M, which has been isolated as a genetic suppressor that stabilizes the structure of UapA in a dimerization defective mutant (Kourkoulou et al., 2019). The L234M mutant has a wild-type like affinity for native substrates, moderately reduced transport rate, and leads to no substrate specificity modification.

**Figure 1.**
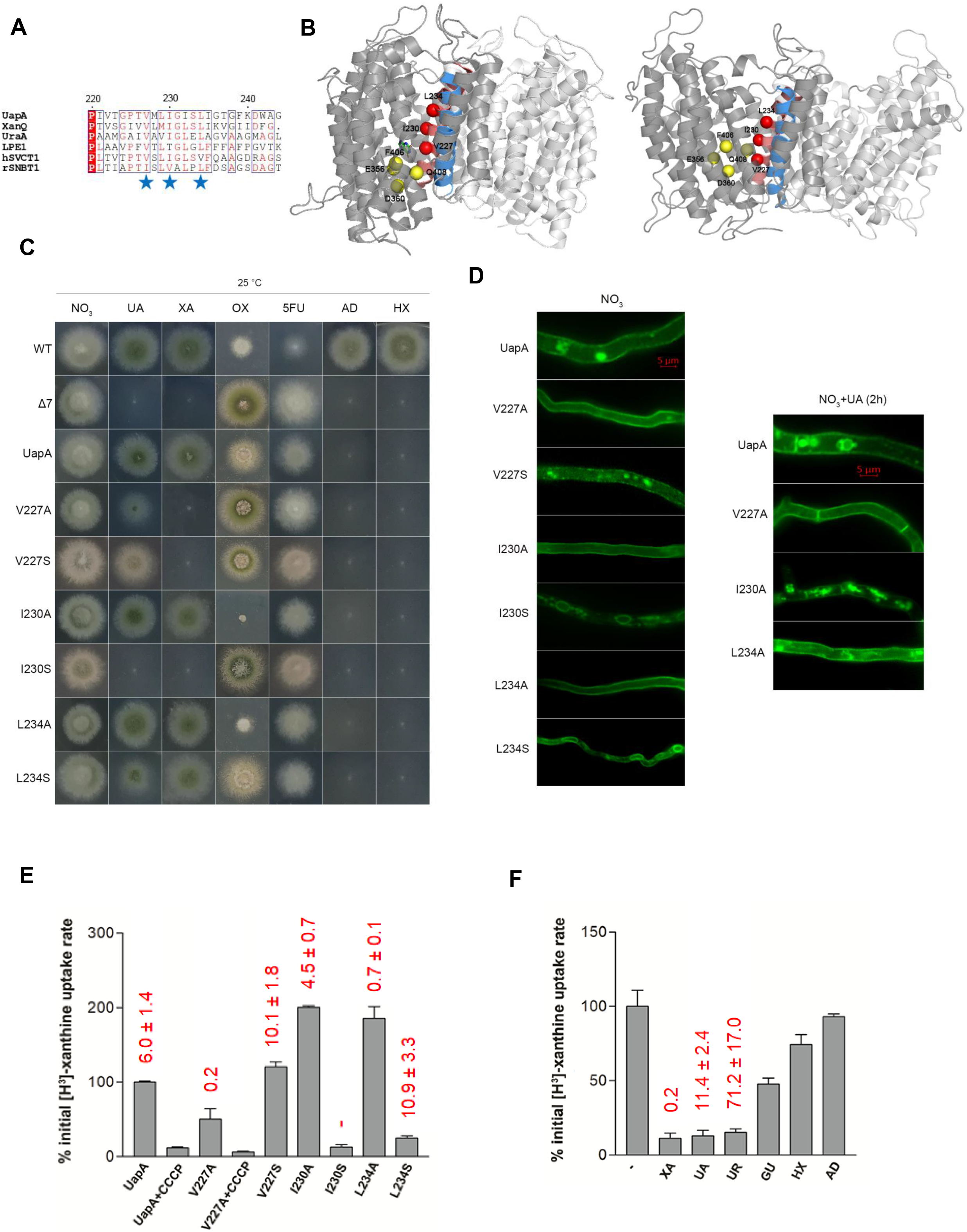
Functional analysis of TMS5 mutants. **(A)** TMS5 sequence alignment of UapA, XanQ, UraA, LPE1, hSVCT1 and rSNBT1. XanQ and UraA are xanthine and uracil transporters respectively in Escherichia coli. LPE1 is a maize xanthine-uric acid transporter. hSVCT1 is a human L-ascorbate transporter. rSNBT1 is a rat purine-uracil transporter (Gournas et al., 2008). Blue stars indicate mutated amino acids. **(B)** Topological models of inward-facing (left) and outward-facing (right) conformations of the UapA dimer highlighting TMS5 (red) and TMS12 (blue) at the interface with the core/elevator domain. Red spheres indicate the positions of the residues mutated (V227, I230, L234). Yellow spheres represent residues in the core/elevator domain, directly involved in substrate binding (E356 and D360 in TMS8 and F406 and Q408 in TMS10). For clarity, annotations are shown in one of the two UapA monomers (deeper grey). Notice the downwards sliding of the core/elevator domain in the inward-facing conformer. **(C)** Growth tests of UapA mutants on minimal media (MM) with purines as N source or nucleobase toxic analogues. UA is uric acid, XA is xanthine, AD is adenine, HX is hypoxanthine, 5FU is 5-flouoruracil, OX is oxypurinol. Growth on 10 mM sodium nitrate (NO_3_^-^) is included as a standard N source control. Purine concentration as N sources is 0.5–2.0 mM. Concentration of toxic analogues is 100 μM. All tests are performed at 25 °C and pH 6.8. Three control strains are included in the growth tests. WT is a wild-type strain possessing all relative endogenous nucleobase-related transporters. Δ7 is a strain lacking all 7 major nucleobase transporters (Krypotou and Diallinas, 2014). UapA is a Δ7 strain expressing wild-type UapA functionally tagged with GFP. All UapA mutant versions shown are expressed and analysed in the Δ7 strain. **(D)** Epifluorescence microscopy showing the subcellular localization of UapA mutants and wild-type UapA in the presence of NO_3_^-^ as N source (left). Notice the perinuclear ER rings in I230S and L234S, and the PM instability (endocytic turnover) in V227S. A similar experiment with wt UapA, V227A, I230A and L234A in the presence of substrate (1mM UA for 2 h) is shown in the right panel. Notice the PM stability of V227A relative to the wild-type UapA and the other mutants. **(E)** Relative ^3^H-xanthine (0.3 μM) transport rates of all mutants analysed expressed as percentages of initial uptake rates (V) compared to the wild-type UapA (UapA) rate. CCCP (Carbonyl cyanide m-chlorophenyl hydrazone) is a proton gradient uncoupler inhibiting wt UapA activity. UapA initial uptake rate is arbitrarily taken as 100%. Km values (μ?) for xanthine, estimated as described in Materials and Methods, are shown at the top of histograms. Results are averages of three measurements for each concentration point. SD was less than 20%. **(F)** Relative H-xanthine (0.3 μM) transport rates of strain V227A in the absence or presence of excess (2 mM) unlabeled nucleobases (UR is uracil and GU is guanine), expressed as percentages of initial uptake rates (V) compared to the rate of V227A in the absence of unlabeled nucleobases, considered as 100%. Km and Ki values (μM) for XA, UA and UR are shown at the top of the histograms. Results are averages of three measurements for each concentration point. SD was less than 20%.

To explore whether residues in TMS5 might affect the dynamics and/or specificity of transport, we constructed and functionally analyzed the following new mutants: V227A, V227S, I230A, I230S, L234A and L234S. The residues modified are located at the interface of the TMS5 helix overlooking the substrate binding site in the sliding elevator domain, and are all extremely conserved in the Nucleobase Ascorbate Transporter (NAT) family (Gournas et al., 2008; Diallinas and Gournas 2008), where UapA belongs (**Figure 1B**). The mutated UapA versions were introduced and analyzed in a genetic background lacking detectable nucleobase-related transport activities due to null mutations in all relevant transporter genes (Δ7), as described in Kourkoulou et al., 2021. *A. nidulans* can grow on purines as N sources and is sensitive to several toxic analogues of nucleobases allowing direct assessment of UapA mutant versions expressed in Δ7 (**Figure 1C**). Mutants I230A and L234A showed normal growth on media with uric acid and xanthine, similarly to an isogenic control strain expressing wild-type (wt) UapA, while L234S showed moderately reduced growth on these purines, and especially on uric acid. I230S scored as a loss-of-function mutation, similar to a strain lacking UapA. Both mutations in V227 led to inability for growth on xanthine and to significantly reduced growth on uric acid, suggesting they might differentially recognize native substrates. We also examined whether the mutations affected UapA-medicated sensitivity to oxypurinol, a toxic purine analogue known to be specifically recognized by UapA. I230A and L234A showed increased sensitivity to oxypurinol compared to the wild-type UapA control, suggesting enhanced activity or/and increased presence of UapA in the PM. This was in sharp contrast to V227A and all Ser-substituted mutants, which showed reduced sensitivity to oxypurinol. Finally, all mutants tested were resistant to 5-FU (toxic analogue of uracil) and could not grow on adenine or hypoxanthine, similar to wild-type UapA (**Figure 1C**).

To test whether the growth phenotypes are related to UapA biogenesis and stability we performed epifluorescence microscopic analysis, taking advantage that all mutants were made in fully functional GFP-tagged UapA (**Figure 1D**). V227A, I230A and L234A all showed proper plasma membrane steady state localization. V227S was localized in the plasma membrane (PM), but also and in cytosolic foci (static vacuoles and motile early endosomes), suggesting intrinsic instability. I230S and L234S showed total or partial ER-retention (i.e. appearance of perinuclear ER rings), respectively, suggesting partial protein misfolding. Stability for the Ala-substituted UapA mutants was also examined in the presence of substrates (e.g. uric acid). It has been shown that UapA undergoes activity-dependent, substrate-elicited, endocytic turnover, similarity to several transporters in fungi (Gournas et al., 2010; Karachaliou et al., 2013). Unlike I230A and L234A, which behaved similar to wild-type UapA, V227A proved highly resistant to substrate-elicited internalization, remaining stably localized in the PM, similar to previously studied loss-of function mutants (Karachaliou et al., 2013). These results showed that UapA biogenesis and PM stability in the absence of substrates are not affected by Ala substitutions in TMS5 (V227A, I230A, L234A). Thus, the UapA-dependent growth phenotypes observed could be directly assigned to UapA activity.

To further understand the effect of the mutations constructed, we performed direct transport assays using radiolabeled xanthine (**Figure 1E**). Mutants I230A and L234A showed increased rate of xanthine transport (2-fold of the wt UapA), in line with growth tests, and stable presence in the PM. Mutants I230S and L234S showed very low transport activities (~10% and 22%, respectively), compatible with growth tests, considering that the threshold for growth on purines as N sources is ~25% of wild-type UapA transport rate. Surprisingly, both V227A and V227S, which scored as loss-of-function mutants in Figures 1C and 1D, showed ‘accumulation’ of radiolabeled xanthine in cells (40 and 115% of wild-type UapA, respectively).

To further investigate the effect of TMS5 mutations on UapA transport function activity, and in particular understand the paradox of V227 substitutions, we measured apparent *K*_m_ values for xanthine (**Figure 1E**). Mutants V227S, I230A and L234S showed moderately affected affinities for xanthine (~4.5-11.0 μM compared to 6 μM in the wt), whereas V227A and L234A showed 30- and 9-fold increased affinities (~0.2-0.7 μM). Thus, while Ser substitutions of V227 or L234 had a moderate effect for xanthine recognition, both Ala substitutions led to a dramatic increase in xanthine binding. In the case of V227A and V227S, the measured binding affinities for xanthine (0.2 and 10.1 μM, respectively) did not justify the lack of UapA-mediated growth on xanthine, where xanthine is provided at ~400 μM. Thus, the simplest explanation of the disagreement of uptake measurements with growth tests is that V227 mutants *bind* tightly, but are unable to *transport*, xanthine. If so, the amount of radiolabeled xanthine detected in uptake assays of V227A or V227S mutants should reflect xanthine ‘trapped’ in UapA molecules present at the PM, rather than *bona fidae* intracellular accumulation of xanthine. Given that V227A has acquired a 30-fold increased binding affinity for xanthine, we also tested whether it shows increased binding affinities for other nucleobases. Uric acid and uracil inhibited the binding radiolabeled xanthine with *K*_i_ values of ~11 and ~71 μM respectively, while guanine, adenine and hypoxanthine showed much lower affinities (*K*_i_ of ~0.5-1.0 mM) **(Figure 1F**). Thus, V227A is a UapA version that recognizes, in some cases with extremely high affinity, but does not transport nucleobase substrates. Specifically in the case of xanthine, dramatically increased binding affinity seems to block transport cycle, which in turn suggests that UapA is trapped in a substrate-occluded conformation. Noticeably also, xanthine apparent trapping in V227A was shown to depend on H^+^ binding, as the use of the CCCP (Carbonyl cyanide m-chlorophenyl hydrazone) proton gradient uncoupler led to dramatic reduction of radiolabeled xanthine associated with cells. Overall, our findings suggest that the character of aliphatic amino acids on the side of TMS5 that faces the sliding-elevator domain is not only critical for proper folding and stability in the PM, as might have been expected, but also for transport dynamics.

### TMS12 mutations

TMS12 is a moderately conserved transmembrane segment in UapA homologues (**Figure 2A**). Previous unbiased genetic screens and rationally designed mutations have shown that specific replacements of little conserved aliphatic amino acids L459, V463 and A469 facing the interface with the sliding elevator domain enlarge the specificity of UapA towards nucleobase substrates (Kosti et al., 2010; Algüel et al., 2016). Here, we constructed and functionally analyzed the following TMS12 mutations: A461G, A461S and I470A. The rationale for mutating A461 was based on a recent mutational analysis of *Escherichia coli* homologues of UapA, and particularly of XanQ, a highly specific xanthine transporter (Tatsaki et al., 2021). In XanQ, residue S377, which corresponds to A461 in UapA and faces the TMS12 side towards the dimerization interface (**Figure 2B**), proved to be an element critical for specificity as its substitution with Gly enlarges the capacity of XanQ to recognize non-native substrates (e.g. guanine). Molecular Dynamics further showed that the S377G replacement tilts TMS12, resulting in a rearrangement of the neighboring F376 relative to F94 (TMS3) and F322 (TMS10) in the substrate binding pocket. These three Phe residues are proposed to pack substrates via pi-pi stacking interactions. Thus, topological modification of this aromatic network might lead to loosening of XanQ specificity. I470 was mutated because it lies at the end of TMS12, at the substrate exit to the cytoplasm (**Figure 2B**).

**Figure 2.**
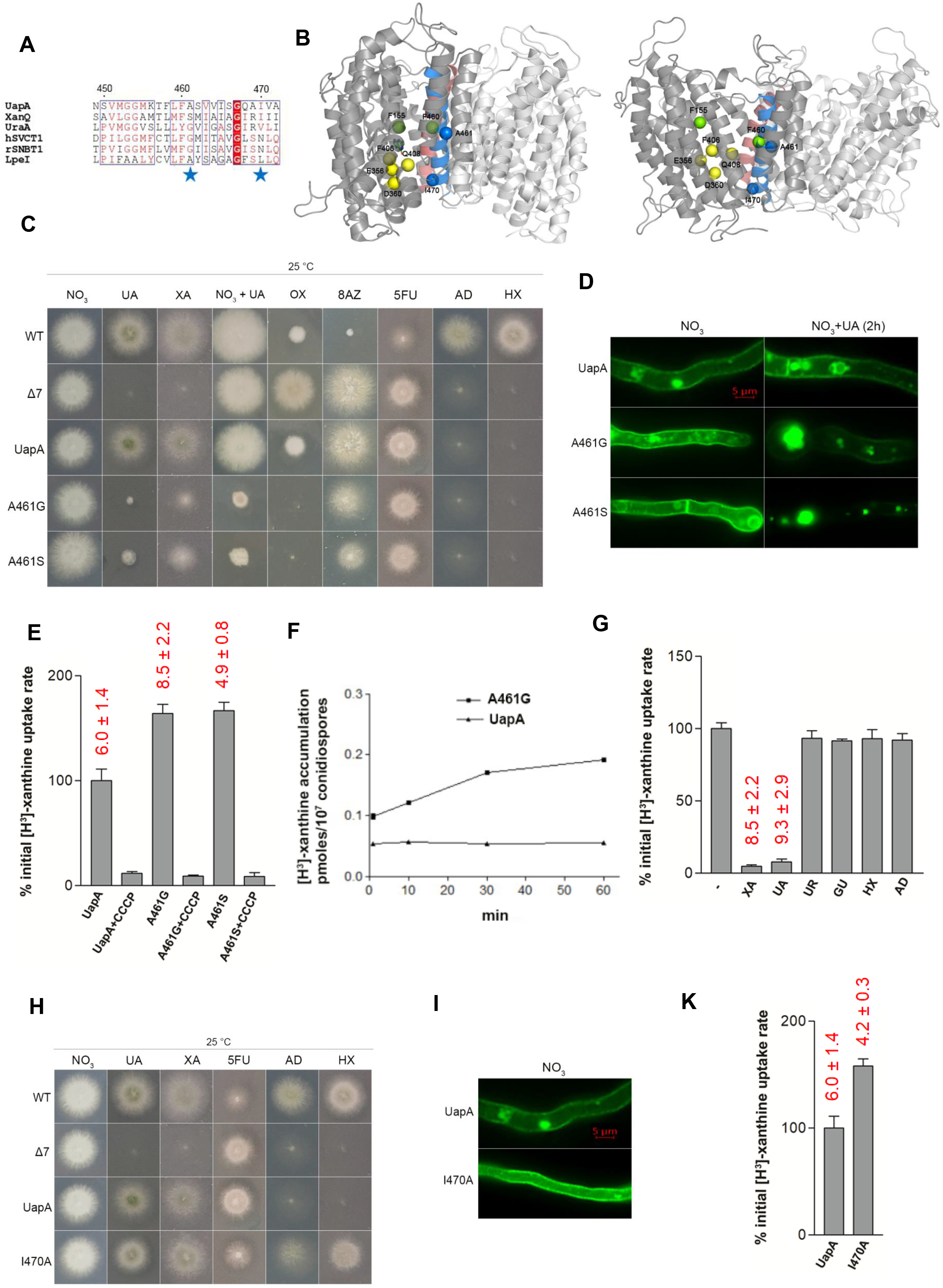
Functional analysis of TMS12 mutants. **(A)** TMS12 sequence alignment of UapA, XanQ, UraA, LPE1, hSVCT1 and rSNBT1. Details on proteins are as in the legend of Figure 1. **(B)** Topological models of inward-facing (left) and outward-facing (right) conformations of the UapA dimer highlighting TMS12 (blue) and TMS5 (red) at the interface with the core/elevator domain. Blue spheres indicate the positions of the residues mutated (A461, I470). Yellow spheres represent residues in the core/elevator domain, directly involved in substrate binding (E356 and D360 in TMS8 and F406 and Q408 in TMS10). Green spheres indicate two Phe (F150 in TMS3 and F460 in TMS12) crucial for the coordination of the substrate to the respective binding site. For clarity, annotations are shown in one of the two UapA monomers (deeper grey). **(C)** Growth tests of UapA mutants A461G, A461S and control strains on minimal media (MM). Details are as described in Figure 1. **(D)** Epifluorescence microscopy showing the subcellular localization of wild-type UapA and UapA mutants (A461G andA461S) in the presence of NO_3_^-^ or NO_3_^-^ + UA (substrate of UapA). **(E)** Relative ^3^H-xanthine (0.3 μM) transport rates of all mutants analysed expressed as percentages of initial uptake rates (*V*) compared to the wild-type UapA (UapA) rate. Details are as in the legend of Figure 1. *K*_m_ values (μM) for xanthine, estimated as described in Materials and Methods, are shown at the top of histograms. Results are averages of three measurements for each concentration point. SD was less than 20%. **(F)** Relative H-xanthine (0.3 μM) transport accumulation in wild-type UapA (control) and A461G strains as a time course. Uptake results are averages of three measurements for each concentration point. SD was less than 20%.**(G)** Relative ^3^H-xanthine (0.3 μM) transport rates of strain A461G in the absence or presence of excess (2 mM) unlabeled nucleobases (UR is uracil and GU is guanine), expressed as percentages of initial uptake rates (*V*) compared to the rate of A461G in the absence of unlabeled nucleobases, considered as 100%. *K*_m_ and *K*_i_ values (μM) for XA, UA and UR are shown at the top of the histograms. Results are averages of three measurements for each concentration point. SD was less than 20%. **(H)** Growth tests of UapA mutant I470A and control strains on minimal media (MM) as described in Figure 1. **(I)** Epifluorescence microscopy showing the subcellular localization of UapA mutant I470A and wild-type UapA in the presence of NO_3_^-^ as N source. **(K)** Relative ^3^H-xanthine (0.3 μM) transport rates of mutant strain I470A expressed as percentage of initial uptake rate (*V*) compared to the wild-type UapA (UapA) rate. UapA initial uptake rate is arbitrarily taken as 100%. *K*_m_ values (μM) for xanthine, estimated as described in Materials and Methods, are shown at the top of histograms. Results are averages of three measurements for each concentration point. SD was less than 20%.

Most interestingly, mutations A461G and A461S led to apparent toxicity of uric acid (**Figure 2C)**. Toxicity can only be explained by considering uncontrolled accumulation of uric acid, a metabolite known to be cytotoxic at high concentrations. A461G and A461S also conferred relative toxicity to xanthine, oxypurinol or 8-azaguanine, the latter being a nucleobase analogue not recognized by wild-type UapA. Otherwise, the A461G and A461S remained resistant to the toxic uracil analogue 5-FU and were incapable of growing on adenine or hypoxanthine, similarly to the control strain expressing wt UapA (**Figure 2C**). Thus, conserved substitutions of A461 seem to modify substrate recognition, but mostly affect UapA transport capacity. Noticeably, mutations A461G and A461S conferred increased instability to the UapA protein, especially in the presence of native substrates (e.g. uric acid), as reflected in increased endocytic turnover (**Figure 2D**). This is often the case with hyperactive mutants that show increased transport rates (Gournas et al., 2010). We confirmed, via direct uptake measurements, that A461 mutations indeed lead to increased, H^+^-dependent, initial transport rate of xanthine, while affecting very little the affinity of UapA for xanthine (**Figure 2E**). Noticeably also, A461G led to significantly increased steady-state accumulation of xanthine, reaching a plateau after 1 h of incubation of radiolabeled xanthine, contrasting the 2 min plateau reached via wild-type UapA activity (**Figure 2F**). We also examined whether A461G affects the binding affinity of uric acid, adenine, hypoxanthine, guanine or uracil, by uptake competition assays. **Figure 2G** shows that the binding profile of A461G in respect to these nucleobases is very similar to that of the wild-type UapA, that is, only uric acid is recognized as a high-affinity ligand/substrate. Overall, the functional profile of A461G and A461S mutants shows that these mutations do not affect substrate binding affinities, but seem to ‘unlock’ UapA transport activity from a control possibly exerted via interactions of TMS12 with residues of the core/elevator domain.

Mutation I470A proved to enlarge UapA specificity to 5-FU and all purines (**Figure 2H**) and did not affect the proper localization and stability of UapA to the PM (**Figure 2I**). Uptake assays further showed that I470 has significantly increased transport rate, while conserving nearly wild-type affinity for xanthine (**Figure 2K**). On the whole, a very similar functional profile has been previously obtained with specific mutations in residues of TMS12 located along the same side of this helix (e.g. mutations in L459, V463 and A469). Thus, specific aliphatic residues located all along the TMS12 side facing the elevator domain control UapA specificity and transport dynamics.

### Epistatic interactions between TMS5 and TMS12

To further understand the role of the two most interesting new mutations, V227A and A461G, which seem to have opposing effects on UapA function (i.e. stabilization of a transport-incompetent occluded conformation due to high-affinity substrate trapping *versus* uncontrolled transport catalysis associated with protein instability), we combined these mutations in a single protein. **Figure 3** reveals that the functional profile of the double V227A/A461G mutant seems to reflect a more or less additive effect of the two single mutations, in respect to transport capacity (**Figure 3A and 3C**), UapA stability (**Figure 3B**), and affinity for xanthine (**Figure 3C**). However, the double V227A/A461G mutant is still kinetically distinct from wild-type UapA, showing higher transport rate and affinity for xanthine.

**Figure 3.**
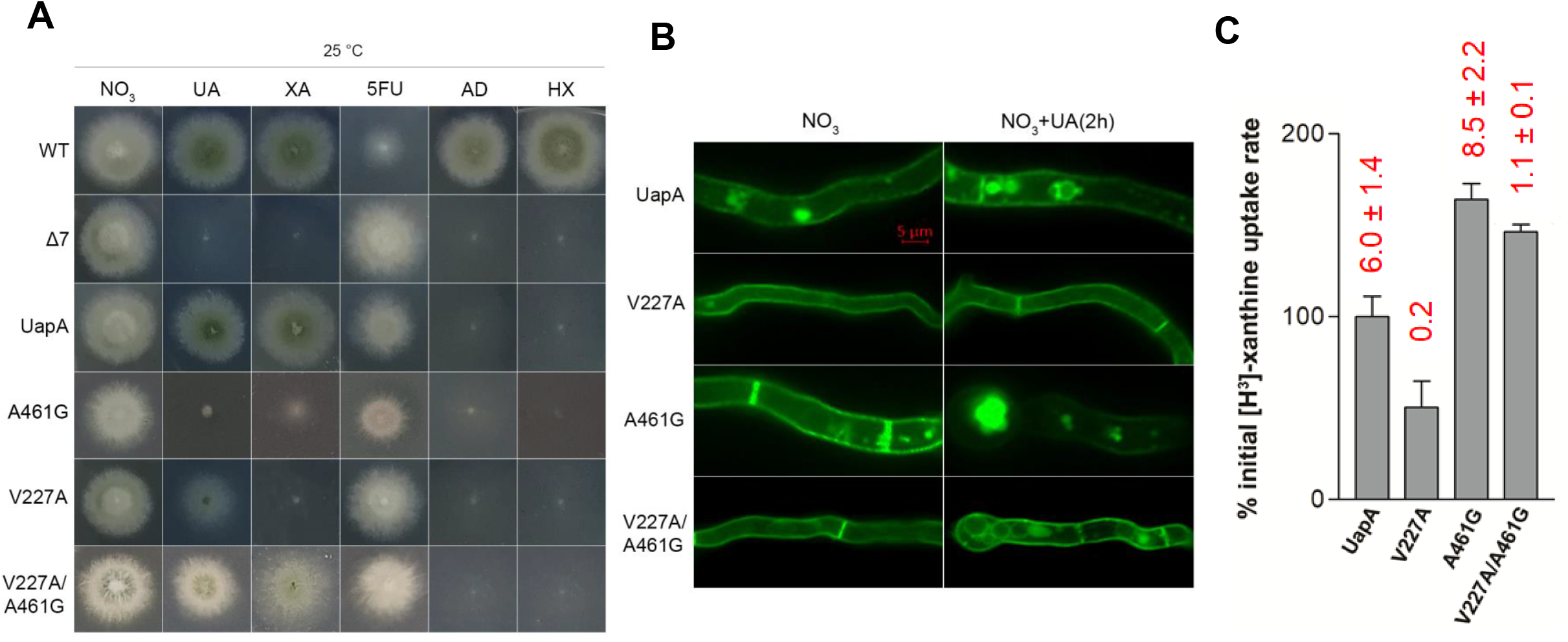
Functional analysis of a strain combining mutations V227A in TMS5 and A461G in TMS12. **(A)** Growth tests of UapA mutants V227A, A461G, V227A/A461G and control strains on minimal media (MM) as described in Figure 1. **(B)** Epifluorescence microscopy showing the subcellular localization of UapA mutants V227A, A461G, V227A/A461G and wild-type UapA in the presence of NO_3_^-^ as N source (left column). Two similar experiments with wild-type UapA, V227A, A461G and V227A/A461G in the presence of substrate (1mM UA for 2 h) are shown in the right column. Notice the increased stability of V227A in contrast to the instability of A461G. V227A/A461G shows an intermediate picture in respect to stability **(C)** Relative H-xanthine (0.3 μM) transport rates of all mutants analysed expressed as percentages of initial uptake rates (*V*) compared to the wild-type UapA (UapA) rate. UapA initial uptake rate is arbitrarily taken as 100%. *K*_m_ values (μM) for xanthine, estimated as described in Materials and Methods, are shown at the top of histograms. Results are averages of three measurements for each concentration point. SD was less than 20%.

### Conclusion

The present work provides new genetic evidence supporting the critical role of TMS5 and TMS12 helices in the dynamics of transport of a sliding elevator-type transporter. Our results show that modifications of transport dynamics leading to ‘substrate-lock’ (i.e. in TMS5) or ‘total unlock’ (i.e. in TMS12) of sliding of the motile elevator domain, carrying the substrate-binding site, can be achieved via specific conservative mutations of aliphatic amino acids. Genetic changes affecting transport dynamics are also shown to affect transporter stability and specificity. This further suggests that in the course of evolution subtle alterations in TMS5 and TMS12 of elevator transporters might have led to transporters with distinct biophysical properties, transport capacities, or specificity. Noticeably, mutations in conserved aliphatic residues in TMS5 affect mostly the dynamics of transport (negatively or positively), while mutations in partially conserved aliphatic residues in TMS12 lead to uncontrolled transport, which in turn converts UapA into a more promiscuous transporter. The epistatic relationships of TMS5 and TMS12 mutations highlight the difficulty in predicting *a priori* the overall function of an elevator transporter based solely on the character of amino acid residues directly interacting with substrates. This work also provides a frame for the genetic stabilization of elevator-type transporters for crystallographic or cryo-EM structural analyses.

## Materials and Methods

### Media, strains and growth conditions

Standard complete (CM) and minimal media (MM) for *A. nidulans* growth were used. Media and supplemented auxotrophies were used at the concentrations given in http://www.fgsc.net (McCluskey et al., 2010). Glucose 1 % (w/v) was used as carbon source. 10 mM sodium nitrate (NO_3_) was used as a standard nitrogen source. Nucleobases, and toxic analogues were used at the following final concentrations: 5-FU, 8-azaguanine and oxypurinol at 100 μM; uric acid, xanthine, adenine, hypoxanthine, guanine, at 0.5-2.0 mM. All media and chemical reagents were obtained from Sigma-Aldrich (Life Science Chemilab SA, Hellas) or AppliChem (Bioline Scientific SA, Hellas). A *ΔfurD::riboB ΔfurA::riboB ΔfcyB::argB ΔazgA ΔuapA ΔuapC::AfpyrG ΔcntA::riboB pabaA1 pantoB100* mutant strain, named Δ7, was the recipient strain in transformations with plasmids carrying *uapA* alleles based on complementation of the pantothenic acid auxotrophy *pantoB100* (Krypotou and Diallinas, 2014). *pabaA1* is a paraminobenzoic acid auxotrophy. *A. nidulans* protoplast isolation and transformation was performed as previously described (Koukaki et al., 2003). Growth tests were performed at 25 ^o^ C for 96 h, at pH 6.8.

### Standard molecular biology manipulations and plasmid construction

Genomic DNA extraction from *A. nidulans* was performed as described in FGSC (http://www.fgsc.net). Plasmids, prepared in *E. coli,* and DNA restriction or PCR fragments were purified from agarose 1% gels with the Nucleospin Plasmid Kit or Nucleospin ExtractII kit, according to the manufacturer’s instructions (Macherey-Nagel, Lab Supplies Scientific SA, Hellas). Standard PCR reactions were performed using KAPATaq DNA polymerase (Kapa Biosystems). PCR products used for cloning, sequencing and re-introduction by transformation in *A. nidulans* were amplified by a high fidelity KAPA HiFi HotStart Ready Mix (Kapa Biosystems) polymerase. DNA sequences were determined by VBC-Genomics (Vienna, Austria). Site directed mutagenesis was carried out according to the instructions accompanying the Quik-Change^®^ Site-Directed Mutagenesis Kit (Agilent Technologies, Stratagene). The principal vector used for most *A. nidulans* mutants is a modified pGEM-T-easy vector carrying a version of the *gpdA* promoter, the *trpC* 3’ termination region and the *panB* selection marker (Krypotou et al., 2015). Mutations were constructed by oligonucleotide-directed mutagenesis or appropriate forward and reverse primers (Table S1).

### Protein Model Construction

The inward-facing UapA model PDB ID is 5i6c. The outward-facing model of UapA was generated using as template the Band3 anion exchanger structural homologue crystallized in the outward conformation, as described in Kourkoulou et al., 2021. The models shown here are presented with PyMOL 2.5 (https://pymol.org).

### Transport assays

Kinetic analysis of wild-type and mutant UapA was measured by estimating uptake rates of [^3^H]-xanthine uptake (40 Ci mmol^-1^, Moravek Biochemicals, CA, USA), as previously described in (Krypotou and Diallinas, 2014). In brief, [^3^H]-xanthine uptake was assayed in *A. nidulans* conidiospores germinating for 4 h at 37° C, at 140 rpm, in liquid MM, pH 6.8. Initial velocities were measured on 10 conidiospores/100 μL by incubation with concentration of 0.2-2.0 μM of [^3^H]-xanthine at 37° C. For the competition experiments, initial uptake rates of [^3^H]-xanthine were measured in the simultaneous presence increasing concentrations (1 μM - 1 mM) of various putative nucleobase-related inhibitors. All transport assays were carried out at least in two independent experiments and the measurements in triplicate. Standard deviation was < 20%. Results were analyzed in GraphPad Prism software.

### Epifluorescence microscopy

Samples for standard epifluorescence microscopy were prepared as previously described (Gournas et al., 2010). In brief, sterile 35 mm l-dishes with a glass bottom (Ibidi, Germany) containing liquid minimal media supplemented with NaNO_3_ and 1% glucose were inoculated from a spore solution and incubated for 16 h at 25°C. The images were obtained using an inverted Zeiss Axio Observer Z1 equipped with an Axio Cam HR R3 camera. Image processing and contrast adjustment were made using the ZEN 2012 software while further processing of the TIFF files was made using Adobe Photoshop CS3 software for brightness adjustment, rotation, alignment and annotation.

## Acknowledgments

We are grateful to Emmanuel Mikros and Iliana Zantza for the outward model of UapA. We thank the lab members Sofia Dimou and Georgia Maria Sagia for critically reading the manuscript.

## Notes

### Competing Interest Statement

The authors have declared no competing interest.

