## Supplementary Table S1 for "Transmembrane helices 5 and 12 control transport dynamics, substrate affinity and specificity in the elevator-type UapA transporter"

### Supplementary Material

| Mutation | Oligonucleotide sequence of forward primer | Oligonucleotide sequence of reverse primer |
|---|---|---|
| **V227A** | CATCGTCACTGGTCCCACTGCAATGCTTATCGGGATAAGTCTGATTGG | CCAATCAGACTTATCCCGATAAGCATTGCAGTGGGACCAGTGACGATG |
| **V227S** | CATCGTCACTGGTCCCACTTCAATGCTTATCGGGATAAGTCTG | CAGACTTATCCCGATAAGCATTGAAGTGGGACCAGTGACGATG |
| **I230A** | GGTCCCACTGTAATGCTTGCCGGGATAAGTCTGATTGG | CCAATCAGACTTATCCCGGCAAGCATTACAGTGGGACC |
| **I230S** | CTGGTCCCACTGTAATGCTTAGCGGGATAAGTCTGATTGGAAC | GTTCCAATCAGACTTATCCCGCTAAGCATTACAGTGGGACCAG |
| **L234A** | GTAATGCTTATCGGGATAAGTGCGATTGGAACTGGGTTCAAAG | CTTTGAACCCAGTTCCAATCGCACTTATCCCGATAAGCATTAC |
| **L234S** | GTAATGCTTATCGGGATAAGTTCGATTGGAACTGGGTTCAAAG | CTTTGAACCCAGTTCCAATCGAACTTATCCCGATAAGCATTAC |
| **A461G** | GGGATGAAGACGTTTCTCTTCGGTTCGGTCGTTATTAGCGGACAG | CTGTCCGCTAATAACGACCGAACCGAAGAGAAACGTCTTCATCCC |
| **A461S** | CGGGATGAAGACGTTTCTCTTCTCTTCGGTCGTTATTAGCGGAC | GTCCGCTAATAACGACCGAAGAGAAGAGAAACGTCTTCATCCCG |
| **I470A** | CGTTATTAGCGGACAGGCGGCAGTGGCCAAGGCGCCGTTC | TTCCGGCGCCTTGGCCACGGCCGCCTGTCCGCTAATAACG |

**Table S1. Oligonucleotides used in this study.** The primers were used for site-directed mutagenesis.
